# Pervasive RNA secondary structure in the genomes of SARS-CoV-2 and other coronaviruses – an endeavour to understand its biological purpose

**DOI:** 10.1101/2020.06.17.155200

**Authors:** P. Simmonds

## Abstract

The ultimate outcome of the COVID-19 pandemic is unknown and is dependent on a complex interplay of its pathogenicity, transmissibility and population immunity. In the current study, SARS coronavirus 2 (SARS-CoV-2) was investigated for the presence of large scale internal RNA base pairing in its genome. This property, termed genome scale ordered RNA structure (GORS) has been previously associated with host persistence in other positive-strand RNA viruses, potentially through its shielding effect on viral RNA recognition in the cell. Genomes of SARS-CoV-2 were remarkably structured, with minimum folding energy differences (MFEDs) of 15%, substantially greater than previously examined viruses such as HCV (MFED 7-9%). High MFED values were shared with all coronavirus genomes analysed created by several hundred consecutive energetically favoured stem-loops throughout the genome. In contrast to replication-association RNA structure, GORS was poorly conserved in the positions and identities of base pairing with other sarbecoviruses – even similarly positioned stem-loops in SARS-CoV-2 and SARS-CoV rarely shared homologous pairings, indicative of more rapid evolutionary change in RNA structure than in the underlying coding sequences. Sites predicted to be base-paired in SARS-CoV-2 showed substantially less sequence diversity than unpaired sites, suggesting that disruption of RNA structure by mutation imposes a fitness cost on the virus which is potentially restrictive to its longer evolution. Although functionally uncharacterised, GORS in SARS-CoV-2 and other coronaviruses represent important elements in their cellular interactions that may contribute to their persistence and transmissibility.

## INTRODUCTION

The emergence of SARS-coronavirus-2 (SARS-CoV-2) in 2019 in Wuhan, China was the start of a worldwide pandemic of frequently severe, fatal respiratory disease termed COVID-19 (1-4). The ultimate outcome of the pandemic in terms of global morbidity will be devastating with a fear that recurrent episodes of COVID-19 disease will occur regularly unless effective medical interventions such as global immunisation can be implemented.

In predicting the future of the COVID-19 pandemic, understanding the ability of a virus to persist at a population level is paramount. Its long term presence is governed by its intrinsic transmissibility and the ongoing existence of susceptible individuals to maintain transmission. Transmissibility in turn depends on factors such as its route of spread, the resilience of the virus in the environment and the duration of host immunity after infection and virus clearance. It additionally crucially depends on host persistence; prolonged shedding of infectious virus enables a larger number of susceptible individuals in contact with an infected host to become infected.

In modelling the spread of SARS-CoV-2, information on many of these factors is becoming available. Of greatest concern, populations, such the UK and the USA which have been severely affected by COVID-19, nevertheless display low levels of population exposure (5-8), indicating that further rounds of infection will not be substantially influenced by herd immunity, even pre-supposing that infection confers long-term protection. Examples from other respiratory coronaviruses in humans (9-11) or enteric coronaviruses in animals (12-14) do not provide much reassurance on the latter. Furthermore, SARS-CoV-2 is highly transmissible through respiratory routes and close contact (15,16), it is relatively stable in the environment (17) and SARS-CoV-2 is shed in substantial amounts from respiratory secretions and is infectious through inhalation and ingestion. The final factor, virus persistence with the infected host and the consequent duration of virus shedding is still poorly documented with a paucity of long-term longitudinal studies of infected individuals so close to the start of the pandemic (see Discussion).

In the current study, the degree of RNA secondary structure within the genomes of SARS-CoV-2 and other human and animal coronaviruses was investigated. This was motivated by our previous observation that human and animal positive stranded RNA viruses capable of virus persistence display a marked, and still largely unexplained association with their possession of structured RNA genomes (18-20). The nature of the folding of genomic RNA exposed in the cytoplasm during replication differs in many respects from that associated with discrete RNA structures with defined functions, such as replication elements and translation initiation. These typically display highly evolutionarily conserved pairings, often with covariant sites, which create specific structures that interact with viral and cellular RNA sequences and proteins. In contrast, genome-scale ordered RNA structure (GORS) in persistent viruses is distributed throughout the genome, and appears agnostic about which specific bases are paired – RNA structures of different HCV genotypes are quite different from each over most of the genome yet the overall degree of folding is relatively constant; structure conservation is only apparent within the 3’ end of NS5B and core gene regions and the untranslated genome termini that have known or suspected replication/ translation functions (20).

Without structural conservation, GORS can be best detected thermodynamically, by comparing the minimum folding energy of a WT sequence with an ensemble of control sequences where the base order of the WT sequence has been shuffled (21,22). As examples, this sequence order dependent structure averages at around 8% in HCV, 9% in foot-and-mouth disease virus and 11% in human pegivirus, similar to the extensively structured ribosomal RNA sequences of animals, plants and prokaryotes (19). The association between possession of GORS and virus persistence extends over all species where information on abilities to persist are documented and has potential predictive value for viruses whose ability to persist is undocumented.

In the current study, we have analysed genomic sequences of SARS-CoV-2 and members of other coronavirus species and genera infecting humans and other mammals for the presence of GORS. The expected and intellectually challenging finding of intense RNA formation in all coronaviruses analysed has been reviewed in the context of what is currently known coronavirus persistence in human and other vertebrate hosts.

## RESULTS

### Detection of GORS in coronavirus genomes

A selection of genome sequences of SARS-CoV-2, SARS-CoV, bat-derived sarbecoviruses were analysed along with representative members of each classified species of coronavirus (listed in Table S1; Suppl. Data). Quantitation of RNA structure formation in each sequence was based upon comparison of minimum free energy (MFE) on folding the native sequence with those of sequence order shuffled controls (a procedure that maintained mono- and dinucleotide frequencies of the native sequence, but otherwise substantially randomised its sequence order). Subtraction of the mean shuffled sequence MFE from the native MFE yielded an MFE difference (MFED) that represent the primary metric for quantifying RNA structure in the current study. SARS-CoV-2, SARS-CoV and bat-derived homologues all showed evidence for large scale RNA structure with mean MFED values of around 15% (Fig. 1; raw data listed in Table S1; Suppl. Data). These were substantially higher than that MFED levels of unstructured viruses (mean value 1.1%) and indeed of the majority of structured +strand RNA viruses displaying host persistence, including HCV (7.5-10.7% and HPgV (12.5%). However, high MFED values were found in all coronaviruses, particularly in several members of the *Betacoronavirus* genus (range 8.6-17.5%), and extremely high in avian virus members of the genus *Deltacoronavirus* (23.4% in Bulbul coronavirus HKU11-934, the highest recorded in all previous analyses of vertebrate RNA viruses).

**FIGURE 1.**
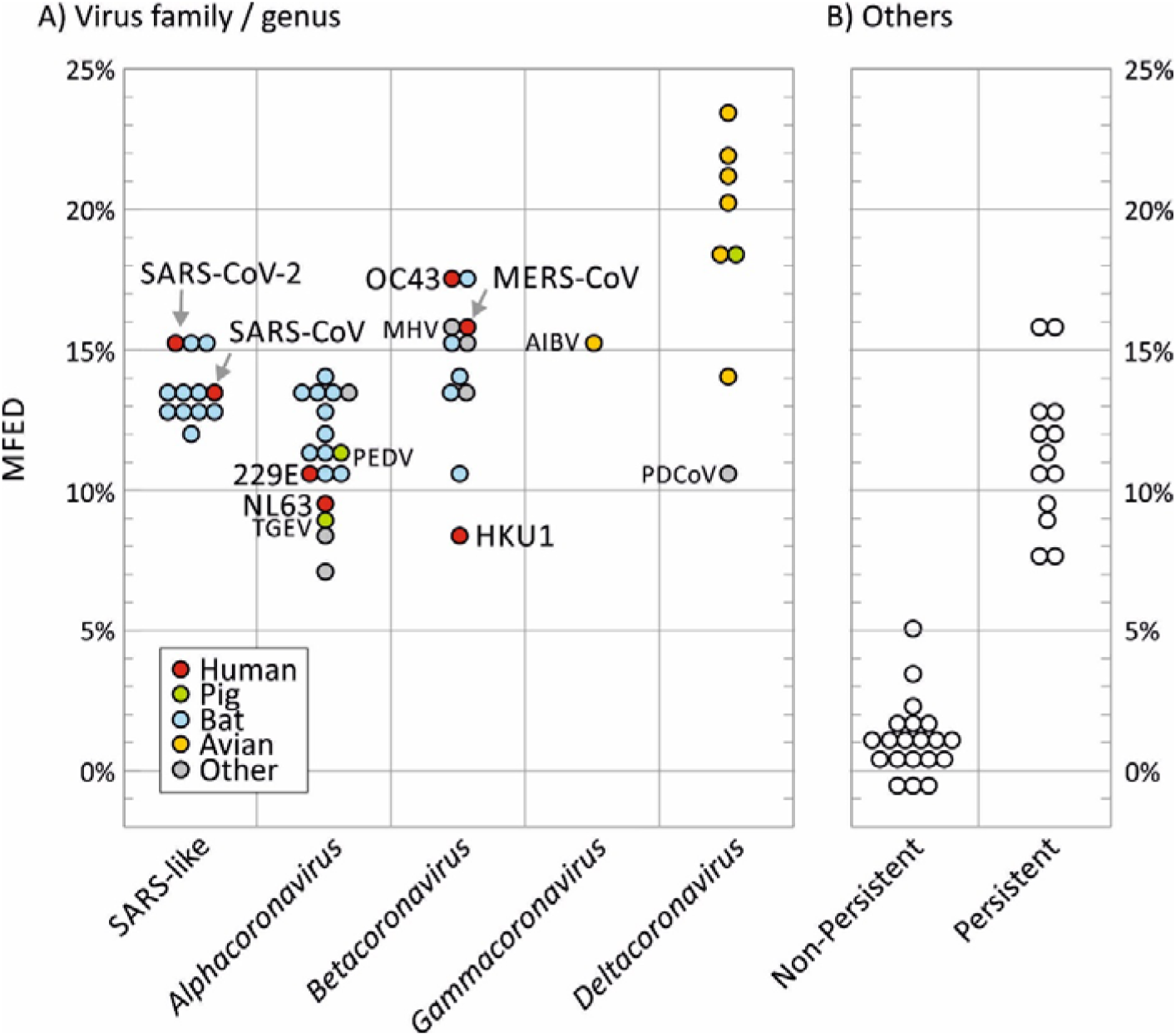
RNA STRUCTURE PREDICTION IN CORONAVIRUSES AND PREVIOUSLY CHARACTERISED PERSISTENT / NON-PERSISTENT +STRAND RNA VIRUSES. RNA structure formation was predicted by comparison of minimum folding energies of virus native sequences with those of shuffled controls (MFED value on the y-axis). (A) Data points represent MFEDs for type member of each currently classified coronavirus species (listed in Table S1; Suppl. Data) and a separate category for SARS-CoV-2, SARS-CoV and a range of SARS-like viruses infecting bats (sarbecoviruses). Human viruses and widely investigated coronaviruses infecting other species are labelled. (B) MFED values of previously analysed +strand mammalian viruses from a previous study and that reported the association between RNA structure and persistence (19).

By analysing MFED values for individual sequence fragments used in MFED calculations, it was apparent that SARS-CoV-2 was structured throughout the genome (Fig. 2). Consistently high values of around 20% were found in the nsp2 and nsp3 genes in the ORF1A-encoding region, around 10-15% in in the remainder of ORF1a and in ORF1b, the spike gene and a peak of >50% in the ORF3a gene. There was no specific association of elevated MFED values and intergenic region, the frameshifting site at the ORF1a/OR1b junction or the 5’ or 3’untranslated regions (UTRs), despite the presence of functional RNA structures in these regions. MFED values in SARS-CoV showed a similar distribution of elevated values as SARS-CoV-2 with some differences in parts of nsp3, spike and ORF3a genes. To investigate the extent to which RNA structure formation imposed constraints on sequence change, variability at synonymous sites in aligned coding sequences of each gene were calculated (green line; Fig. 2). SARS-CoV-2 and SARS-CoV are genetically distinct from each other throughout the genome, but low values indicating constraints did not associate closely with high MFED values or vice versa.

**FIGURE 1.**
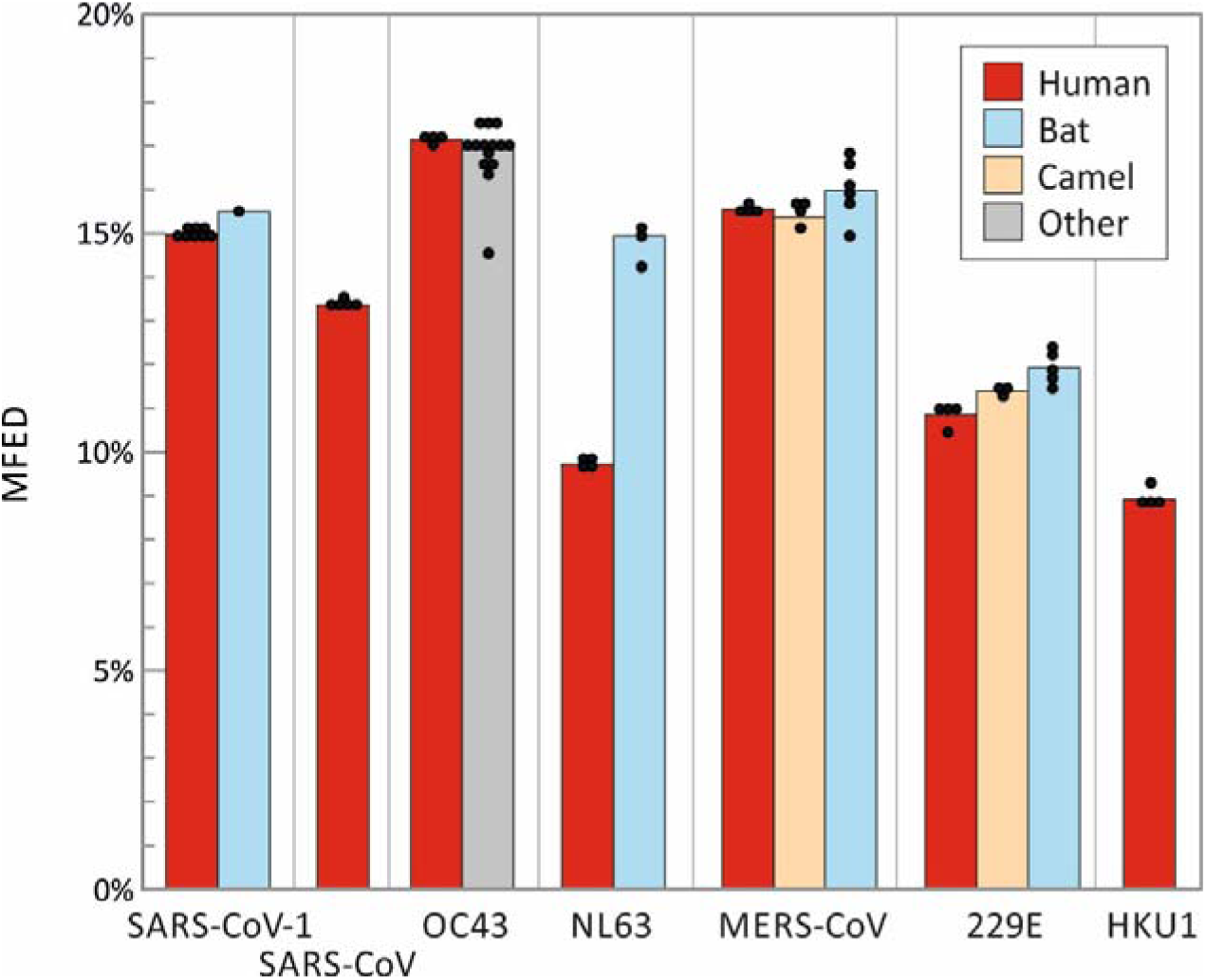
MFED VALUES OF HUMAN CORONAVIRUSES AND THEIR CLOSEST HOMOLOGUES IN OTHER HOST SPECIES. Mean MFED values for selections of representative sequences of each the seven human coronaviruses and their closest homologues in other mammalian species considered to be their zoonotic source. Sequence selection was limited to up to four for each species listed in Table S2; Suppl. Data and displayed as individual points. Significance tests were not attempted as sequences were phylogenetically related.

Each of the human seasonal coronavirus has a known or suspected zoonotic origin (reviewed in (23)), with closely related homologues of OC43 identified in cows, NL63, 229E and MERS-CoV in bats. SARS-CoV-2 is closely related to a coronavirus identified in a bat species (2) that may also represent its ultimate zoonotic source. No genetically close homologues of SARS-CoV or HKU1 are known. Each homologue showed MFED score similar to those of human viruses, although all four bat virus groups were invariably marginally more structured than their human counterparts (SARS-CoV-2, NL63, MERS-CoV and 229E). However, the significance of these differences in difficult to evaluate statistically as the members of each group are phylogenetically related and MFED values derived for individual virus strains do not constitute independent observations.

### Analysis of coronavirus RNA secondary structures

The genomes of SARS-CoV-2 and other coronaviruses are large and visualisation of their genome-wide RNA structure elements by conventional RNA drawings is problematic. I recently developed a contour plotting method for depicting the positions and variability of secondary structure elements in alignments of virus sequences (20). In this method, pairing predictions from RNAFOLD are recursively scanned for stem-loops and unpaired bases in terminal loops of each identified and assigned a height of zero on the z-axis, with genome position and sequence number recorded in the x- and y-axes in a 3–dimensional plot (Fig. 4A). Paired bases either side of the terminal loop were successively plotted according to a colour scale that reflects their distance in the stem relative to the terminal loop. The resulting plot therefore provides an approximate visualisation of the positions, shapes and sizes of RNA structure elements across whole alignments. The 3-dimensional representation can be transformed to a 2-dimensional plot with height indicated by colour coding (Fig. 4B).

**FIGURE 3.**
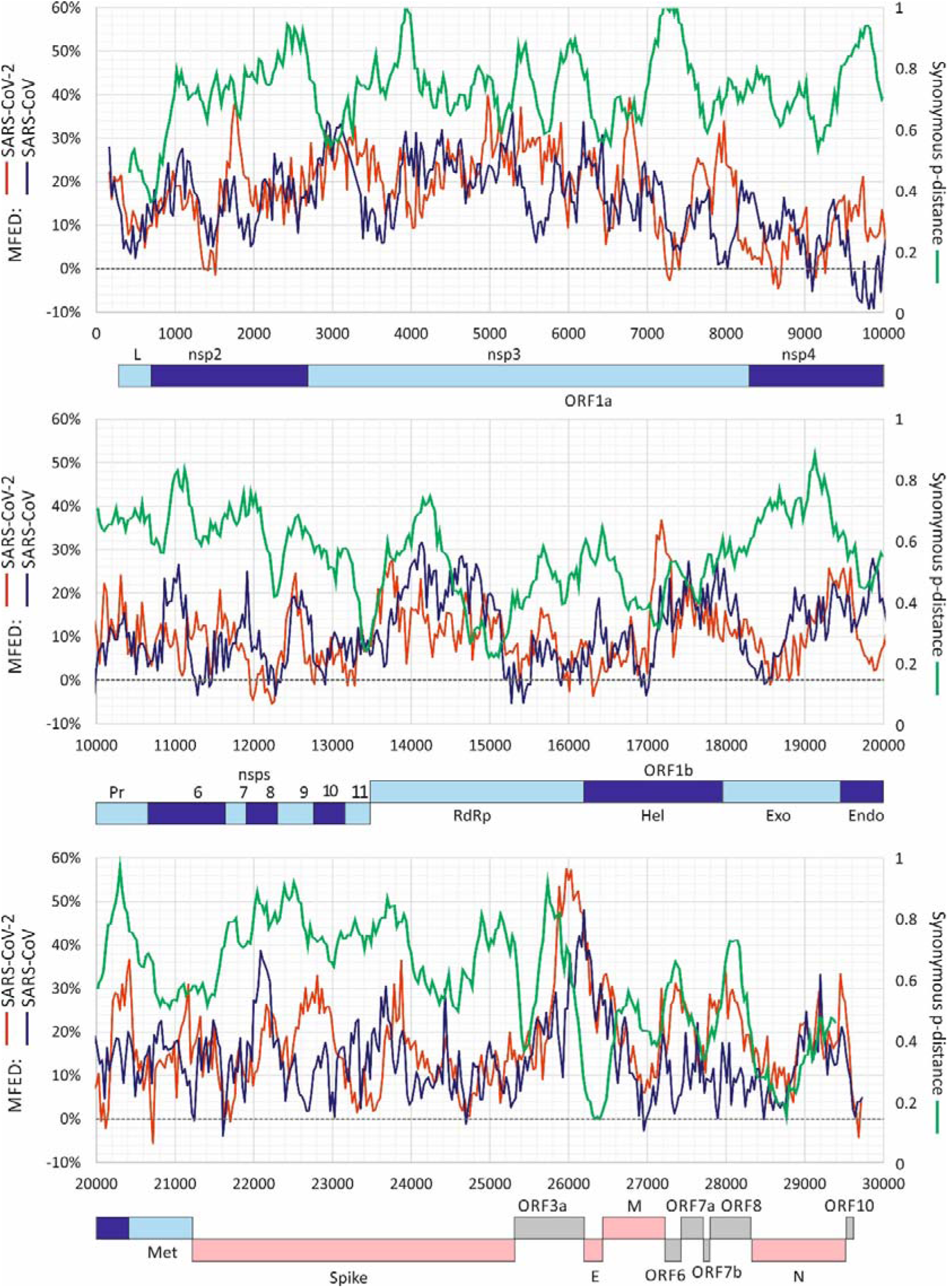
GENOME SCAN OF FOLDING ENERGIES AND SYNONYMOUS VARIABILITY. Windowed MFED values of SARS-CoV-2 and SARS-CoV across the genome (left y-axis) using a fragment size of 350 bases incrementing by 30 bases between fragments. A windowed scan of synonymous p-distances (sequential 300 base fragments incrementing by 30 bases between fragments) of aligned concatenated coding region sequences between SARS-CoV-2 and SARS-CoV is superimposed. A genome diagram of SARS-CoV-2 is drawn to scale under each graph. A listing of the sequences analysed in provided in Table S3; Suppl. Data.

**FIGURE 4.**
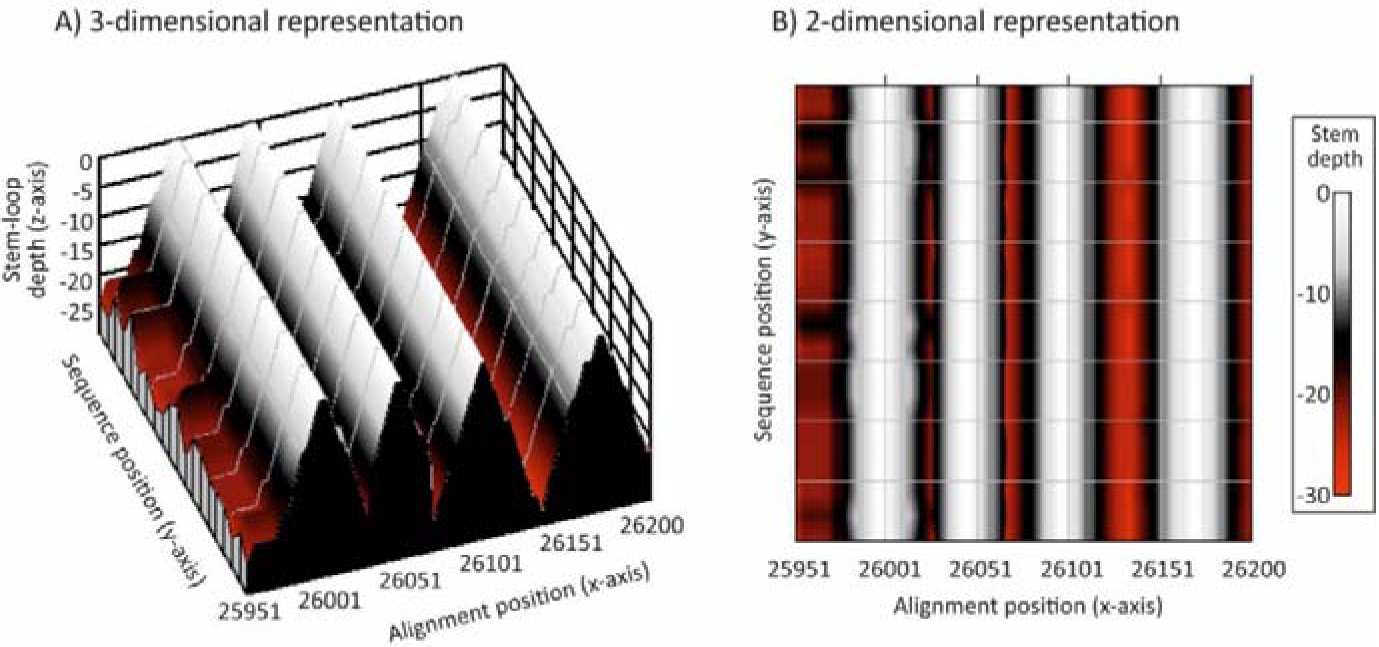
REPRESENTATION OF RNA SECONDARY STRUCTURE AS COUNTOUR PLOTS. Visualisation of RNA structure by contour plotting. Predicted consensus positions of terminal loops in assigned depths of zero, and depths based on numbers of sequential pairings in duplex regions plotted on the z-axis as (A) depth in a 3-dimensional plot and (B) as colour coding. The predicted RNA structure corresponds to a short region of the ORF-3a gene of SARS-CoV-2 analysed in Figs. 5, 6 and 7.

A contour plot was made of an alignment of SARS-CoV-2, SARS-CoV and bat-derived sarbecoviruses (Fig. 5). Variants within SARS-CoV-2 and SARS-CoV were minimally divergent and each produced essentially the same structure predictions. However, these were somewhat different from each other and from bat sarbecoviruses throughout large parts of the genome, highlighting regions with quite different RNA secondary structural organisation of duplex and unpaired regions. Larger scale analyses of two regions of the SARS-CoV-2 and SARS-CoV genomes (positions 2601 – 3400 [in ORF1a] and 25601 – 26400 [in ORF3a / E]) were performed (Fig. 6) to highlight the similarities and differences in base pairings between viruses. Both regions corresponded to areas of high MFED values, 24.5% and 22.32% for SARS-CoV-2 and SARS-CoV in the ORF1a, and 35.8% and 24.7% in ORF3a / E. In the ORF1a region, stem-loop predictions were markedly different between the two viruses despite both viruses showing high MFED values and indeed a consistent pattern of elevation across the entire ORF1a/1b gene, despite these and consistently different actual pairings between the two viruses (Fig. 5).

**FIGURE 5.**
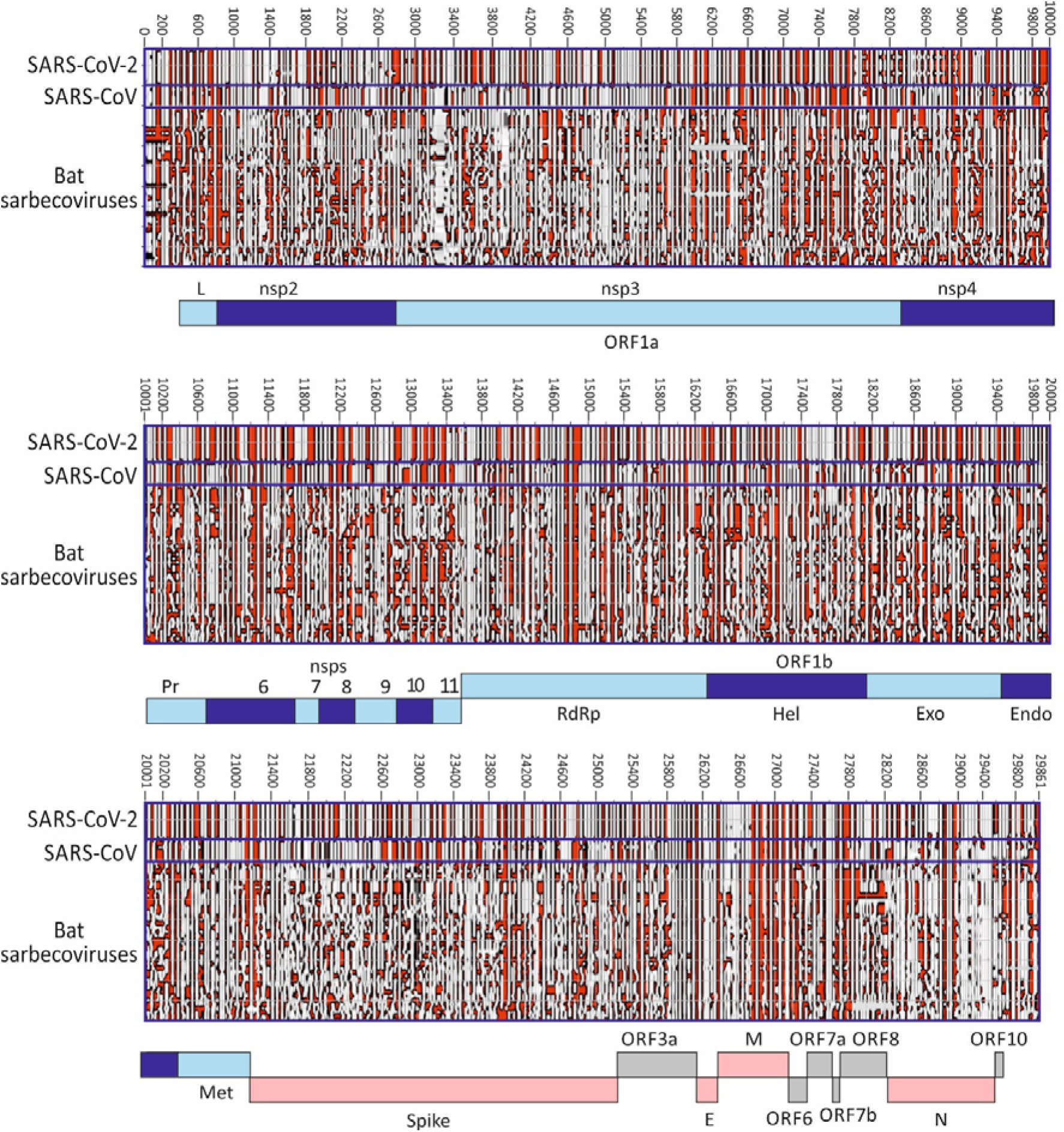
CONTOUR PLOTS OF SARS-CoV-2. SARS-CoV AND BAT SARBECOVIRUSES. Representation of RNA structure elements in the whole genomes of a selection of SARS-CoV-2 (n=9) and other sarbecoviruses (labelled on x-axis; listed in Table S3; Suppl. Data) using the previously described contour plotting method (20).

**FIGURE 6.**
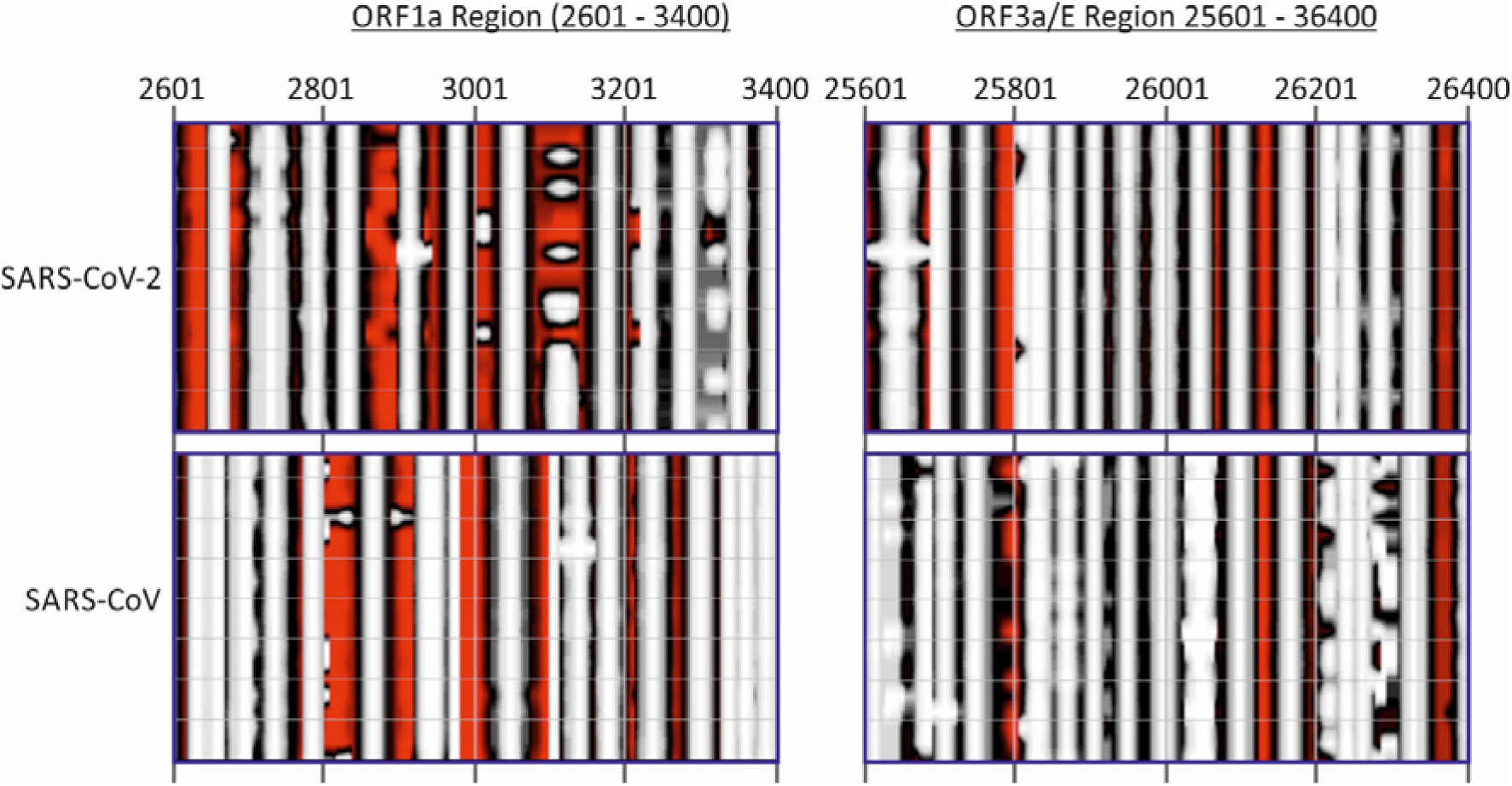
CONTOUR PLOT COMPARISON OF TWO REGIONS OF HIGH MFED VALUES FOR SARS-CoV-2 AND SARS-CoV

In the NS3a / E region, a greater degree of RNA structure conservation was evident in the contour plot. Most predicted stem-loops located to the same places in the alignment, although on closer examination of the base identities of the duplex regions, the actual pairings were non-homologous in the majority of stem-loops (grey dotted arrows in Fig. 6). Despite alignment of the sequences by nucleotide and amino acid sequence identity (and conservation with other sarbecoviruses), duplexes were often formed by distinct bases in the two viruses. For example, pairings in the first stem-loop in SARS-CoV-2 were displaced 5’ by 2 nucleotide positions in the corresponding SARS-CoV sequence (−2). Pairing displacements of -3 (SL4), -7 (SL8), +3 (SL9), -5 (SL10), +6 (SL12) and -16 (SL13) were observed in otherwise similarly positioned and shaped secondary structure elements, with only SL2 and SL5-SL7 showing evidence for homologous pairing. These observations, recapitulated to even greater extents throughout the remainder of the genome, indicate a considerably faster evolution of RNA secondary structure than their underlying coding sequences. For comparison, RNA structures in OC43 and a set of homologues from animals (pigs, cows, camels, giraffe, deer and dogs) were visualised in a separate contour plot (Fig. S1; Suppl. Data). This similarly depicted widely distributed stem-loops through the genome and a degree of structure conservation consistent with the lower degree of sequence divergence between the variants analysed.

Secondary structure elements is SARS-CoV-2 and other coronaviruses primarily comprised simple, sequential stem-loop with single unpaired terminal unpaired regions. A total of 657 stem-loops were predicted for SARS-CoV-2, comparable to totals for other coronaviruses (range 500-625), formed from 2015 duplex regions of 3 or more consecutive base pairs (Table S4; Suppl. Data). Duplexes in stem-loops were frequently interrupted to avoid paired regions longer than 14 consecutive base-pairs. The length distributions of duplex regions were similarly comparable between different coronaviruses (Fig. S2; Suppl. Data).

### Influence of RNA secondary structure on viral diversity

While the functional basis for the adoption of pervasive RNA secondary structure is unknown, the apparent requirement for extensive base-pairing in SARS-CoV-2 and other coronaviruses genomes would be expected to impose constraints on sequence change. Most individual mutations in paired sites would have the effect of weakening RNA secondary structures and lead to a greater phenotypic cost than changes at unpaired sites. For all coronaviruses analysed, approximately 62-67% of bases were predicted be paired (Table S4; Suppl. Data), and their pairing constraints may lead to a substantial restriction on sequence diversification.

To investigate this, sites in an alignment of 17,518 sequences of SARS-CoV-2 were catalogued for diversity through generating a list of the number of sequence changes at each nucleotide site. The terminal 200 bases at each of the genome were excluded from the analysis because of lower coverage and greater frequency of sequencing errors in these regions. Overall, a total of 7064 of the 26468 nucleotide positions analysed were polymorphic (27%). Of the variable sites, approximately one half were represented in two or more sequences (sequence divergence ≥0.0002), declining steeply thereafter (Fig. S5; Suppl. Data). Site variability was compared with predictions of whether they were base paired or non-base paired using RNAFOLD (Fig. 8). The normalised proportion of unpaired and paired sites was similar for sites showing single mutations (variability 0.001), but there was increasing over-representation of unpaired bases at sites showing greater sequence divergence (nearly 2-fold for sites with variability greater than 0.008). This over-representation was even more marked for C-U transitions (blue bars; up to 3.5-fold over-representation). These observations provide evidence for the restricting effect of base-pairing on mutations becoming fixed in the genome.

**FIGURE 7.**
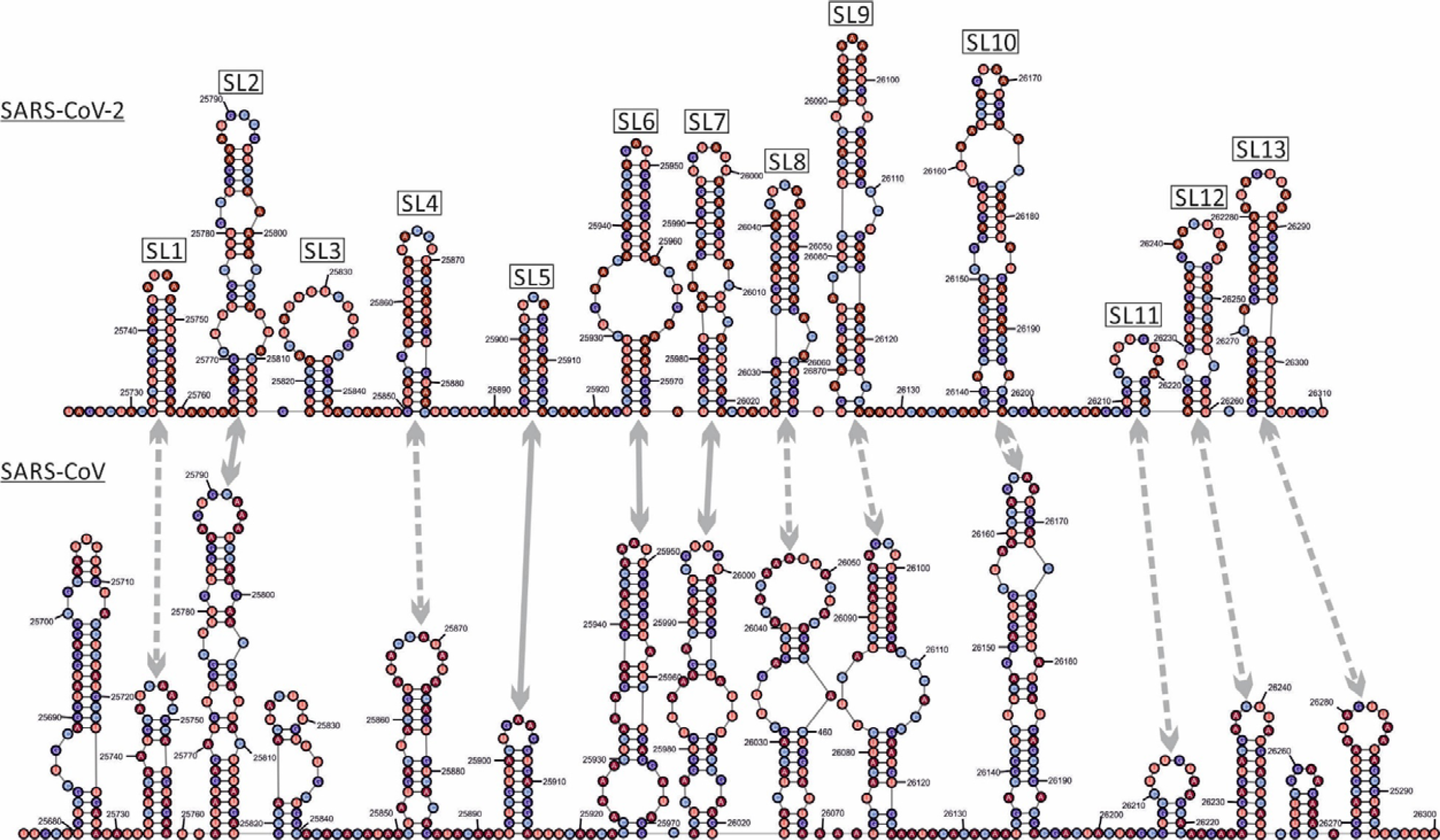
SECONDARY STRUCTURE PREDICTIONS IN THE NS3a / E REGION GENOMIC REGION OF SARS-CoV-2 AND SARS-CoV. Drawing of the predicted RNA secondary structure pairings of genome fragments from 25601-26400 of SARS-CoV-2 and an aligned region of SARS-CoV (24.9% pairwise divergence). Homologous stem-loops between the structure predictions are arrived; sold line: similar structure and homologous pairings; dotted arrow: structures containing non-homologous pairings.

**FIGURE 8.**
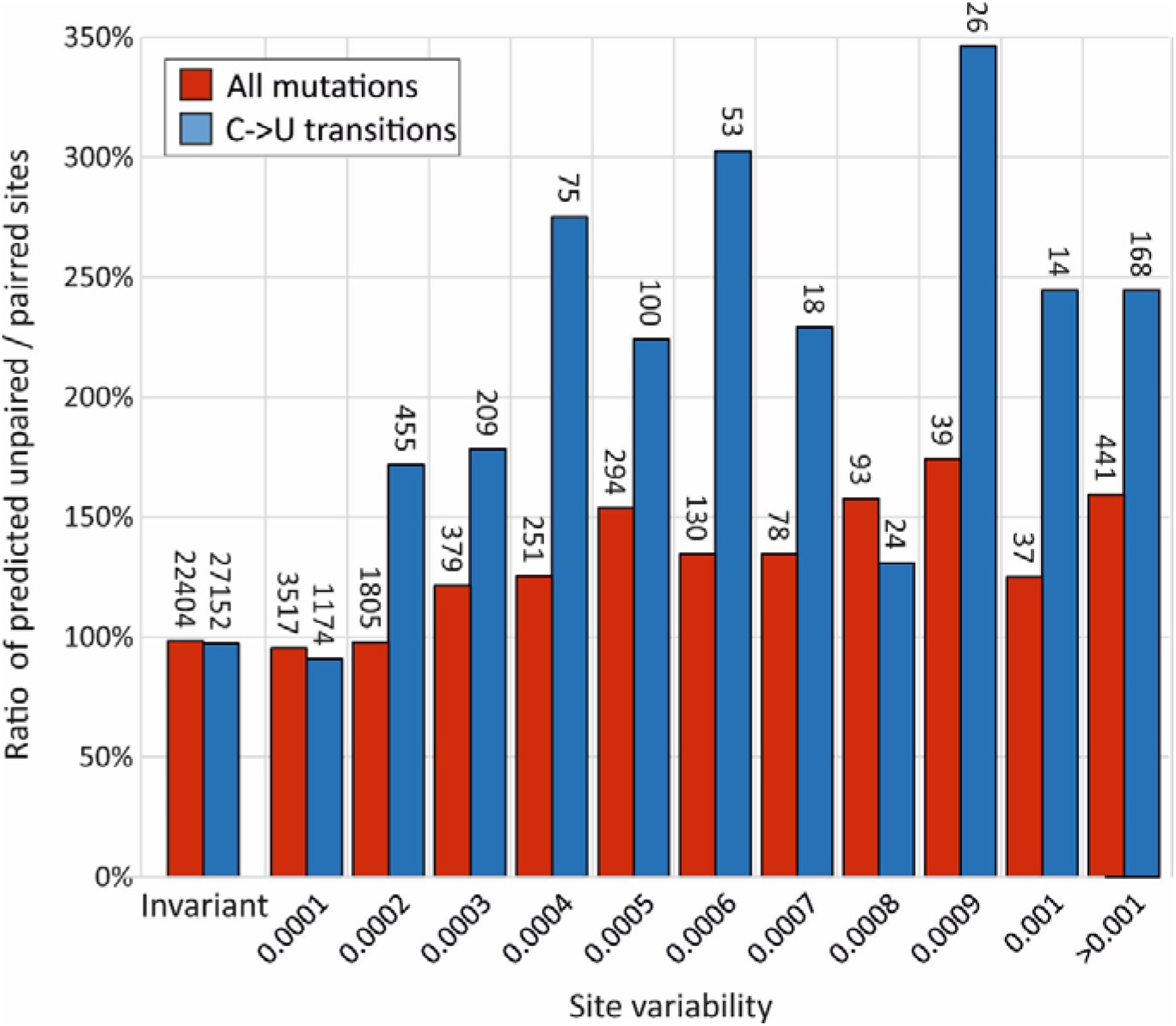
INFLUENCE OF BASE-PAIRING ON SEQUENCE VARIABILITY. Ratios of unpaired to paired sites predicted by RNAFOLD at invariant sites (left two columns) and those showing different degrees of site variability. The numbers of sites in each category are shown above the bars. C>U transitions were the most frequent mutations observed in the dataset and showed a greater influence of base-pairing on their occurrence.

## DISCUSSION

### Prediction of RNA secondary structure

The primary evidence for the existence of RNA structure formation in SARS-CoV-2 and other coronavirus genomes derived from high MFED values across the genome. Values of 15% in SARS-CoV-2, 17% in OC43 (and up to 24% in a deltacoronavirus) are unprecedentedly high compared to those documented for HCV (7-9%, HPgV-1 (11%) and a range of others reported to possess genome scale ordered RNA structure (18,19). MFED calculations identify the sequence order contribution to RNA folding, where elevated values arising from folding energies of native sequences being greater than those of shuffled controls. The use of the NDR shuffling algorithm (24) that preserves these mononucleotide ad dinucleotide compositional features, including the unusual underrepresentation of C and overrepresentation of U in most coronavirus sequences (25,26) provides reassurance that the folding energy differences represent the effects of biologically conditioned sequence ordering to create or maintain RNA secondary structure in coronavirus genomes. Recently published findings of elevated MFEs and consistent outlier Z scores (27) correspond to what are calculated as MFED values in the current study are consistent with conclusion reached about the genome-wide nature of RNA formation.

An independent method to detect and characterise RNA folding, including the identities of individual base-pairings, is based on the detection of covariance. Evidence for pairings is provided by observation of compensatory changes to maintain binding. In this respect, the extremely limited variability of SARS-CoV-2, SARS-CoV, MERS-CoV and indeed of each of the sequence datasets of seasonal coronaviruses prevented this approach from being usefully applied. A second problem is that large scale RNA structure in other viruses, such as HCV, is not necessarily conserved in the same way as it might be in functional RNA structure elements (20). We recently documented substantial variability in pairing sites both between HCV subtypes in large areas of the genome, with structure conservation restricted to functionally mapped *cis-*acting replication elements in the NS5B region and in stem-loops of undefined function in the core gene (28-33). Covariance detection therefore cannot be applied to verify pairing sites in viruses with GORS. Evidence for an analogous lack of pairing constraints and comparably rapid evolution of RNA structure is provided by comparison of RNA structure predictions for SARS-CoV-2 and SARS-CoV (Figs. 5-7). While there is some similarity in the positions and sizes of predicted stem-loop across their genomes (Fig. 5), particularly apparent in the ORF3a / E region (Fig. 6), the actual pairings forming shared stem-loop were non-homologous with frequent displacement of paired based between viruses even though the sizes and spacings of stem-loops were often quite conserved (Fig. 7). This form of “extended” or “inexact” covariance is apparent throughout the SARS-CoV-2 and SARS-CoV genome and supports the idea that it is simple maintenance of pairing rather than functional properties of the stem-loops that are formed that is driving RNA structure formation in coronavirus genomes.

This conclusion is supported by the sheer scale of RNA structure in the SARS-CoV-2 genome. This possesses perhaps 650 or more separate stem-loops across the genome – accepting that many of these predicted structures may derive simply from “over-folding” by energy minimisation programs such as RNAFOLD, even half that number would be far too numerous to plausibly possess specific replication functions. Furthermore, areas of high MFED values did not associate with gene boundaries where discrete RNA structure elements may participate in mRNA processing, frameshifting or other replication functions (34,35), many elements of which have been recently mapped in the SARS-CoV-2 genome (27,36). A similar disconnect between MFED values and functional RNA structures in HCV has been described previously (20). As proposed, it appears that it is the folding of RNA, rather than the structures formed, that drive the creation of GORS; how this modifies interactions of the replicating virus with the cell is discussed below.

### Evolutionary constraints of RNA secondary structure

Notwithstanding the potential inaccuracies of a proportion of specific pairing predictions made by RNAFOLD, the marked difference in sequence variability at paired and unpaired sites (Fig. 8) provides evidence that pairing requirements influence SARS-CoV-2 adaptive fitness and potentially limit its longer-term evolutionary trajectory. A striking observation was the frequency dependent over-representation of variability at unpaired sites; sites showing only single sequence mutations were equally well represented predicted paired and unpaired sites, while those showing multiple changes were substantially over-represented.

The current SARS-CoV-2 datasets maintained by GenBank and GISAID are well curated and consensus sequences generated by NGS methods, particularly with high read depths, rarely contain sequencing errors. However, even a very low frequency of technical mis-assignments in a sequence dataset of over 17,000 full genome sequences will inevitably contain errors and these may have contributed to the lack of association with pairing. Nevertheless, a further and potentially more significant contributor to the large number of single sequence mutations (n=3517) may be the sporadic occurrence of mutations occurring in founder viruses infecting individuals that possess minor fitness defects. These may prevent their propagation and inheritance in other SARS-CoV-2 strains and lack of representation in multiple sequences in the larger dataset. The observation that multiply represented and evolutionarily successful mutations were 2-3 times more likely to occur at unpaired sites indicates that disruption of RNA base-pairing imposes a substantial phenotypic penalty on SARS-CoV-2.

Of the 12 possible mutations, C->U transitions were the most commonly observed in the dataset, consistent with their previously proposed origin through specific RNA editing events by APOBEC or related cytidine deaminases (25,37). Transitions induced by C->U changes were more influenced by pairing constraints than other mutations with nearly 3-fold more occurring at unpaired sites in multiply represented sites. This over-representation and their consequent greater likelihood of inheritance or appearing convergently implies a reduced fitness cost that associated with other mutations. The fact that a substitution of a C for a U at a paired site with G will nevertheless maintain pairing albeit with a lower pairing strength is consistent with this model. The only other mutation that could maintain pairing, A->G, was relatively rare but showed a similar over-representation in variable unpaired sites (141%); however insufficient numbers of mutations occurred for formal frequency analysis (data not shown).

Collectively, the analysis provides evidence that base-pairing imposes a substantial constraint on the diversification of SARS-CoV-2 and presumably of other coronaviruses with comparable degrees of RNA structure formation.

### Biological effects of large-scale RNA structure in SARS-CoV-2 and other coronaviruses

Despite the description of GORS in HCV and a range of other positive-strand RNA viruses, little is known about the biological effects of large-scale RNA structure in viral genomes and how it may influence interactions with the cell. Biophsysically, structured genomes take on a globular, compacted appearance on atomic force microscopy and sequences are inaccessible to external probe hybridisation (19), indicating a quite different RNA configuration from unstructured viruses and potentially influencing interactions with the cell. Maintenance of RNA structure is costly in evolutionary terms, since most changes at paired sites, and potentially a proportion at unpaired sites, disrupt RNA folding. In a previous bioinformatic experiment, 5% simulated evolutionary drift of HCV, HPgV-1 and FMDV reduced MFED values of each virus genome by >50% (18). In the actual world, longer term sequence change in these viruses occurs under constraints to maintain a relatively fixed level of internal base-pairing. The observation that SARS-CoV-2 site diversity was substantially influenced by its predicted pairing (Fig. 8) provides a further indication of the potential phenotypic costs of RNA structure disruption.

A further uncertainty about the purpose and mechanisms of GORS-associated structures is the as yet unexplained correlation between RNA structure formation and virus persistence (18,19). Among many possibilities, we have previously suggested that decreased virus recognition by the innate immune system may fail to activate interferon and other cytokine secretion from infected cells, leading to downstream defects in macrophage and T cell recruitment and maturation. These ultimately may blunt adaptive immune responses sufficiently to enable virus persistence. The poor T helper functions associated with proliferation defects and deletions of reactive CD4 lymphocytes cell responses in those with persistent infections (38-40). Downstream impairment of CD8 cytotoxic T cell and antibody responses may originate from this failure of immune maturation.

On the face of it, the finding that not only SARS-CoV-2, but also all four of the seasonal human coronaviruses possess intensely structured genomes does not square with the previously noted association of GORS with persistence. The human seasonal coronaviruses are considered to cause transient and most often inapparent or mildly symptomatic respiratory infection, notwithstanding the dearth of focussed studies on durations of virus shedding and potential sites of replication outside the respiratory tract. Interestingly, repeat testing of individuals with diagnosed NL63, OC43 and 229E infections within 2-3 months revealed frequent occurrences of infections with the same virus, >20% in the case of NL63 (9). In many cases infections were by the same clade of virus and often showed higher viral loads than observed in the original time point. These findings were interpreted as evidence for reinfection as described in previous studies (10,11), and for some individuals, intermediate samples were obtained and shown to be PCR negative. However, the findings do not rule out persistence over the 3 months of the sampling interval. The observation of NL63 detection in 21% of follow up samples in a study group where only 1.3% of individuals were initially infected provides some tentative support for the latter possibility. Even if the result of re-infection, the findings demonstrate that seasonal coronaviruses fail to induce any effective form of protective immunity from re-infection even over the short period after primary infection. This resembles findings for HCV, where a potentially comparable immunological defect leads to those who have cleared infection to be readily re-infected with same HCV genotype (41,42).

In non-human hosts, coronavirus infections are typically persistent where investigated. These include bovine coronavirus (BCoV) which establishes long-term, asymptomatic respiratory and enteric infections in cows (43,44). BCoV is closely related to OC43 in humans and potentially its zoonotic source (23). Although not longitudinally sampled, MERS-CoV was detected at frequencies >40% in several groups of dromedary camels, similarly indicative of persistence (45) despite its more frequent clearance in infected humans (46). Other coronaviruses showing long term persistence include mouse hepatitis virus, feline calicivirus (47) and infectious bronchitis virus in birds (48,49). Pigs are infected with a range of different coronaviruses of variable propensities to establish persistent infections (50-53). Many of the coronaviruses characterised in pigs have arisen in major outbreaks potentially from zoonotic sources, including porcine deltacoronavirus in 2014 from a sparrow CoV, and porcine epidemic diarrhea virus in 1971 and swine acute diarrhoea syndrome-coronavirus in 2016 from bats (reviewed in (54)). A lack of host adaptation immediately after recent zoonotic spread may contribute the varying outcomes of pig coronavirus infections. Coronaviruses in bats are distributed in the *Alpha-* and *Betacoronavirus* genera, widespread and highly genetically diverse and host specific. Establishing whether infections are persistent in bats is problematic in a standard field study setting. However, high detection rates in faecal samples from bats, including 26% and 24% in large samples of *Minopterus australis* and *M. schreibersii* in Australia (55), 29% in rhinolophid bats in Japan (56) and 30% in various bat species in the Philippines (57) are strongly indicative of persistence. Overall, coronaviruses clearly have a propensity to persist although their ability to achieve this may depend on their degree of host adaptation.

Turning to recently emerged coronaviruses in humans, the course of SARS-CoV infections can be prolonged, up to 126 days in faecal samples (58) although little information on persistence was collected before the outbreak terminated. MERS-CoV infections are persistent in camels but show variable outcomes in humans with respiratory detection and faecal excretion typically ceasing 3-4 weeks after infection onset (59,60) but with individual case reports of much longer persistence in some individuals (46). Based on what is known for other coronaviruses, SARS-CoV-2 clearly has the potential for persistence and indeed probably is persistent in its immediate bat source, *Rhinolophus affinis* (2). Its current presentation as an acute, primarily respiratory infection may represent the typical course of a recently zoonotically transmitted virus with the potential for future adaptive changes to increases its systemic spread and achieve a degree of host persistence apparent in many animal coronaviruses.

Even in the current pandemic period 6 months after the zoonotic event, relatively long periods of respiratory sample detection and faecal excretion of SARS-CoV-2 have been documented, in many cases of greater than 1 month duration (61-65). These occur in both mild and severe cases of COVID-19 in patients, and without co-morbidities or evident immune deficits that may separately contribute to persistence. While the world anxiously awaits how SARS-CoV-2 transmissibility and pathogenicity may evolve in future outbreaks, understanding the mechanisms of post-zoonotic adaptation of SARS-CoV-2 to humans is of crucial importance. Interactions of SARS-CoV-2 with innate immune pathways potentially modulated by large-scale RNA structure may represent one element in this adaptive process.

## MATERIALS AND METHODS

### SARS-CoV-2 and other coronavirus datasets

Coronavirus sequences analysed in the study were downloaded from Genbank. A full listing of their accession numbers and download dates provided in Table S1 (Suppl. Data).

### RNA structure prediction

MFED values were calculated by comparing minimum folding energies (MFEDs) for WT and sequences shuffled in order by the algorithm NDR. For analysis, coronoavirus sequences were split into 350 base sequential sequence fragments incrementing by 15 bases between fragments. For each, MFEs were determined using the RNAFold.exe program in the RNAFold package, version 2.4.2 (66) with default parameters. Summary MFED values (Fig. 1, 2) were based on mean MFEDs for all fragments in the coding regions of each virus sequence. MFED scans were based on averaging MFEDs from sequence sets for each fragment and plotting values out on the y-axis, using the midpoint fragment position on the x-axis (Fig. 3). All shuffling, MFE and MFED determinations were automated in the program MFED scan in the SSE v1.4 package (24)(http://www.virus-evolution.org/Downloads/Software/).

Contour plots were produced as previously described (20). Briefly, ensemble RNA structure predictions were made from sequential 1600 base fragments of the alignment incrementing by 400 bases between fragments using the program SubOpt.exe in the RNAFold package. Those supported by >50% of sub-optimal structures were for constructing a consensus contour plot. SubOpt.exe was invoked through the program StructureDist in the SSE package. A listing of paired and unpaired sites was obtained from the Pos.Dat output from StructureDist. Statistics on stem-loop numbers, duplex and terminal loop lengths was obtained from the Stats List.DT1 file generated by the same program.

### Other analysis

Calculation of synonymous pairwise distances and lists of sequence changes at each site were generated by the programs Sequence Distances, Sequence Changes and Sequence Join in the SSE package. RNA structure drawing were generated from output from Structure Editor in the RNAstructure package (http://rna.urmc.rochester.edu/RNAstructure.html). Statistical analysis and construction of frequency histograms used SPSS version 2

## ACKNOWLEDGEMENTS

The work was supported by a Wellcome Investigator Award Grant WT103767MA.

