## Supplementary Data for "Pervasive RNA secondary structure in the genomes of SARS-CoV-2 and other coronaviruses – an endeavour to understand its biological purpose"

SUPPLEMENTARY FIGURE 1 – CONTOUR PLOT OF OC43 AND HOMOLOGUES IN ANIMALS

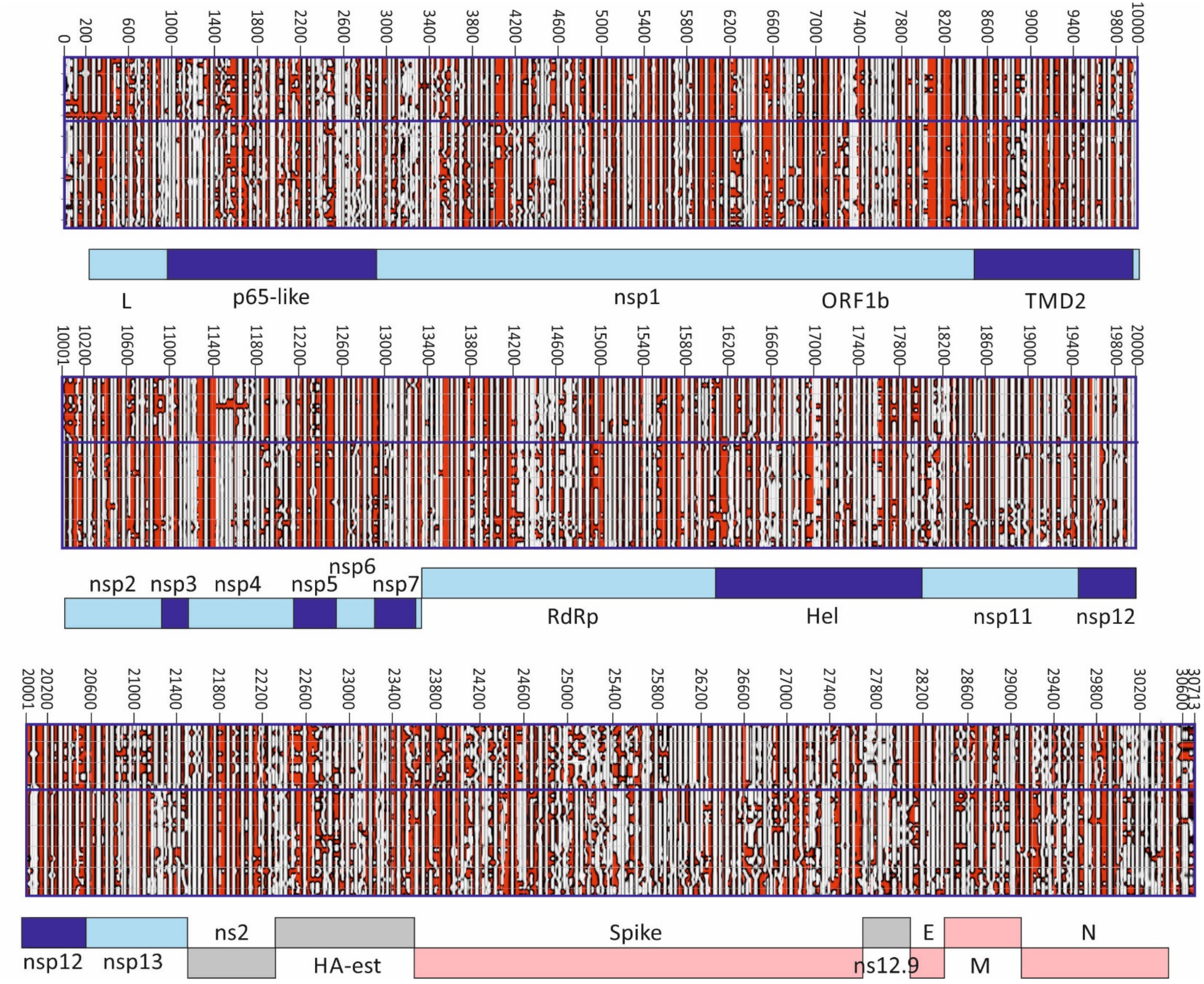

Human OC43 strains (upper panel) and a set of homologues from animals (pigs, cows, camels, giraffe, deer and dogs; lower panel) were aligned with a genome representation of OC43 strain AY585228, using the annotation provided.

FIGURE S2

LENGTH DISTRIBUTION AND POSITIONS OF STEM-LOOP DUPLEXES IN CORONAVIRUSES

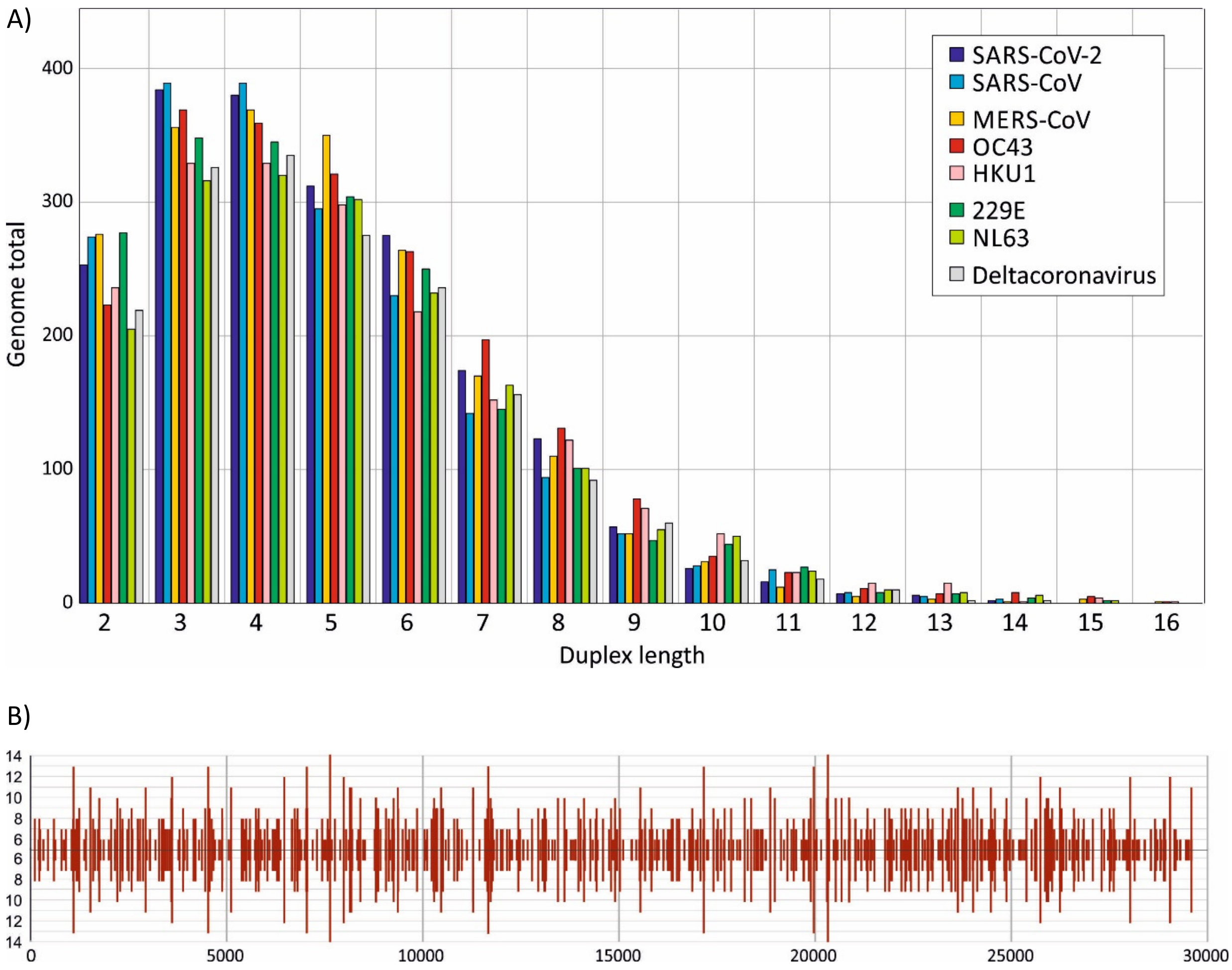

A) Length distribution of uninterrupted duplexes in predicted RNA secondary structures of coronaviruses. (B) Analysis of pairing predictions from the SARS-CoV-2 genome showing the positions and lengths of stem-loop duplexes of length greater than 5 base pairs; the maximum duplex length detected was 14 (n=2).

FIGURE S3

NUMBERS OF VARIABLE SITES IN THE SARS-CoV-2 GENOME

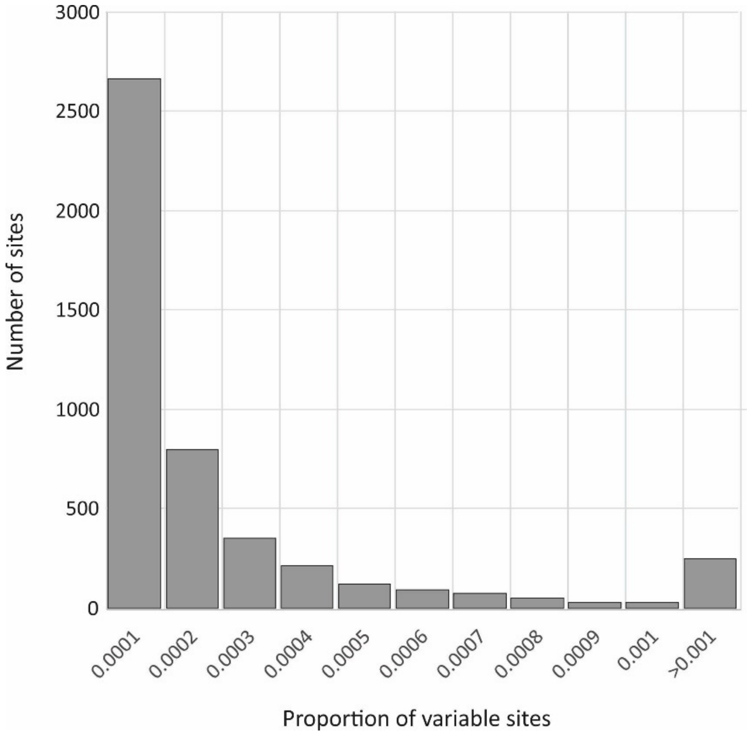

Numbers of sites showing different degrees of sequence variability in a total of 13562 SARS-CoV-2 genomes

TABLE S1

REPRESENTATIVE CORONAVIRUSES USED FOR RNA STRUCTURE ANALYSIS

| Group | Accession_no | Isolate | MFED |
|---|---|---|---|
| *Sarbecovirus** | MN988713 | Severe acute respiratory syndrome coronavirus 2 | 15.29% |
| *Sarbecovirus** | MN996532 | Bat coronavirus RaTG13 | 15.41% |
| *Sarbecovirus** | FJ882953 | SARS coronavirus MA15 ExoN1 isolate P3pp4 | 13.48% |
| *Sarbecovirus** | KF294457 | SARS-related bat coronavirus isolate Longquan-140 | 12.80% |
| *Sarbecovirus** | KJ473813 | BtRf-BetaCoV/SX2013 | 11.92% |
| *Sarbecovirus** | KP886809 | Bat SARS-like coronavirus YNLF_34C | 13.23% |
| *Sarbecovirus** | KJ473814 | BtRs-BetaCoV/HuB2013 | 13.68% |
| *Sarbecovirus** | MG772934 | Bat SARS-like coronavirus isolate bat-SL-CoVZXC21 | 15.51% |
| *Sarbecovirus** | KY352407 | Severe acute respiratory syndrome-related coronavirus | 12.77% |
| *Sarbecovirus** | GU190215 | Bat coronavirus BM48-31/BGR/2008 | 12.75% |
| *Sarbecovirus** | JX993988 | Bat coronavirus Cp/Yunnan2011 | 13.57% |
| *Sarbecovirus** | MK211374 | Coronavirus BtRl-BetaCoV/SC2018 | 12.82% |
| *Alphacoronavirus* | KF430219 | Bat coronavirus CDPHE15/USA/2006 | 13.35% |
| *Alphacoronavirus* | JQ989270 | Rousettus bat coronavirus HKU10 isolate 183A | 11.40% |
| *Alphacoronavirus* | KJ473807 | BtRf-AlphaCoV/HuB2013 | 11.42% |
| *Alphacoronavirus* | AF304460 | Human coronavirus 229E | 10.37% |
| *Alphacoronavirus* | KF294380 | Lucheng Rn rat coronavirus isolate Lucheng-19 | 13.35% |
| *Alphacoronavirus* | LC119077 | Ferret coronavirus FRCoV4370 | 7.35% |
| *Alphacoronavirus* | HM245925 | Mink coronavirus strain WD1127 | 8.14% |
| *Alphacoronavirus* | EU420138 | Miniopterus bat coronavirus 1 | 13.37% |
| *Alphacoronavirus* | EU420139 | Bat coronavirus HKU8 strain AFCD77 | 13.89% |
| *Alphacoronavirus* | KJ473806 | BtMr-AlphaCoV/SAX2011 | 13.06% |
| *Alphacoronavirus* | KJ473809 | BtNv-AlphaCoV/SC2013 | 10.86% |
| *Alphacoronavirus* | AF353511 | Porcine epidemic diarrhea virus strain CV777 | 11.14% |
| *Alphacoronavirus* | DQ648858 | Bat coronavirus (BtCoV/512/2005) | 11.87% |
| *Alphacoronavirus* | EF203064 | Bat coronavirus HKU2 strain HKU2/GD/430/2006 | 10.63% |
| *Alphacoronavirus* | AY567487 | Human Coronavirus NL63 | 9.65% |
| *Alphacoronavirus* | KY073745 | NL63-related bat coronavirus strain BtKYNL63-9b | 13.73% |
| *Alphacoronavirus* | AJ271965 | Transmissible gastroenteritis virus | 8.75% |
| *Betacoronavirus* | AY585228 | Human coronavirus OC43 strain ATCC VR-759 | 17.57% |
| *Betacoronavirus* | KM349742 | Betacoronavirus HKU24 strain HKU24-R05005I | 15.22% |
| *Betacoronavirus* | AY597011 | Human coronavirus HKU1 genotype A | 8.60% |
| *Betacoronavirus* | AY700211 | Murine hepatitis virus strain A59 | 15.55% |
| *Betacoronavirus* | KF636752 | Bat Hp-betacoronavirus/Zhejiang2013 | 15.26% |
| *Betacoronavirus* | KC545383 | Betacoronavirus Erinaceus/VMC/DEU/2012 | 13.60% |
| *Betacoronavirus* | EF065509 | Bat coronavirus HKU5-1 | 13.50% |
| *Betacoronavirus* | EF065505 | Bat coronavirus HKU4-1 | 17.50% |
| *Betacoronavirus* | KU762338 | Rousettus bat coronavirus isolate GCCDC1 356 | 10.86% |
| *Betacoronavirus* | EF065513 | Bat coronavirus HKU9-1 | 14.13% |

| | | | |
|---|---|---|---|
| *Gammacoronavirus* | IBACGB | Avian infectious bronchitis virus | 14.98% |
| *Deltacoronavirus* | JQ065048 | Wigeon coronavirus HKU20 strain HKU20-9243 | 14.21% |
| *Deltacoronavirus* | FJ376619 | Bulbul coronavirus HKU11-934 | 23.45% |
| *Deltacoronavirus* | JQ065043 | Porcine coronavirus HKU15 strain HKU15-155 | 18.30% |
| *Deltacoronavirus* | FJ376622 | Munia coronavirus HKU13-3514 | 20.24% |
| *Deltacoronavirus* | JQ065044 | White-eye coronavirus HKU16 strain HKU16-6847 | 21.92% |
| *Deltacoronavirus* | JQ065047 | Night-heron coronavirus HKU19 strain HKU19-6918 | 18.50% |
| *Deltacoronavirus* | JQ065049 | Common-moorhen coronavirus HKU21 strain HKU21-8295 | 21.19% |
| *Deltacoronavirus* | EU111742 | Coronavirus SW1 | 10.71% |

TABLE S2

CORONAVIRUS SEQUENCES USED FOR MFED COMPARISON IN DIFFERENT HOSTS

| **Run_name** | **Sequence** | **Host** | **MFED** |
|---|---|---|---|
| SARS-CoV-2 | MN988713 | Human | 15.05% |
| SARS-CoV-2 | MT093571 | Human | 14.85% |
| SARS-CoV-2 | MT049951 | Human | 14.98% |
| SARS-CoV-2 | MT039890 | Human | 14.99% |
| SARS-CoV-2 | MT027064 | Human | 14.92% |
| SARS-CoV-2 | MT007544 | Human | 15.04% |
| SARS-CoV-2 | MN994467 | Human | 15.10% |
| SARS-CoV-2 | MN996528 | Human | 15.01% |
| SARS-CoV-2 | MN996527 | Human | 14.99% |
| Sarbecovirus* | MN996532 | Bat | 15.07% |
| SARS-CoV-1 | FJ882953 | Human | 13.33% |
| SARS-CoV-1 | AY654624 | Human | 13.33% |
| SARS-CoV-1 | FJ882926 | Human | 13.45% |
| SARS-CoV-1 | FJ882943 | Human | 13.27% |
| SARS-CoV-1 | HQ890531 | Human | 13.35% |
| NL63_H | AY567487 | Human | 9.59% |
| NL63_H | KY674916 | Human | 9.81% |
| NL63_H | JQ765564 | Human | 9.78% |
| NL63_H | MG428700 | Human | 9.73% |
| NL63_B | KY073744 | Bat | 15.14% |
| NL63_B | NC_048216 | Bat | 14.22% |
| NL63_B | KY073746 | Bat | 14.97% |
| HKU1_H | AY597011 | Human | 8.91% |
| HKU1_H | KF686342 | Human | 8.85% |
| HKU1_H | DQ415899 | Human | 9.28% |
| HKU1_H | KY674921 | Human | 8.75% |
| OC43_H | AY585228 | Human | 17.19% |
| OC43_H | KY369907 | Human | 17.03% |
| OC43_H | KF530088 | Human | 17.15% |
| OC43_H | KF530060 | Human | 17.23% |
| OC43_OM | KU558922 | Bovine | 16.63% |
| OC43_OM | MG757140 | Bovine | 16.57% |
| OC43_OM | KY419105 | Pig | 16.33% |
| OC43_OM | EF424622 | Camel | 17.45% |
| OC43_OM | EF424623 | Camel | 17.57% |
| OC43_OM | EF424624 | Camel | 17.53% |
| OC43_OM | FJ425186 | Deer | 17.08% |
| OC43_OM | MG518518 | Deer | 16.96% |
| OC43_OM | MH810163 | Deer | 14.53% |
| OC43_OM | KF906249 | Camel | 17.02% |
| OC43_OM | MN514964 | Camel | 17.05% |

| | | | |
|---|---|---|---|
| OC43_OM | MN514962 | Camel | 16.96% |
| OC43_OM | JX860640 | Dog | 16.93% |
| OC43_OM | KX432213 | Dog | 16.75% |
| 229E_H | AF304460 | Human | 10.44% |
| 229E_H | KY967357 | Human | 11.04% |
| 229E_H | KY996417 | Human | 10.90% |
| 229E_H | JX503060 | Human | 10.93% |
| 229E_C | KT368905 | Camel | 11.49% |
| 229E_C | MF593473 | Camel | 11.43% |
| 229E_C | JQ410000 | Camel | 11.34% |
| 229E_B | KT253272 | Bat | 11.50% |
| 229E_B | KY073747 | Bat | 11.91% |
| 229E_B | KT253269 | Bat | 11.64% |
| 229E_B | KY073748 | Bat | 12.16% |
| 229E_B | MN611517 | Bat | 12.38% |
| MERS-CoV_H | KC164505 | human | 15.57% |
| MERS-CoV_H | KT026454 | human | 15.59% |
| MERS-CoV_H | KT156561 | human | 15.49% |
| MERS-CoV_H | KM027255 | human | 15.55% |
| MERS_CoV_C | MH734115 | Camel | 15.67% |
| MERS_CoV_C | MF598699 | Camel | 15.18% |
| MERS_CoV_C | MF598619 | Camel | 15.73% |
| MERS_CoV_C | MG923479 | Camel | 15.41% |
| MERS_CoV_B | MF593268 | Bat | 16.89% |
| MERS_CoV_B | KC869678 | Bat | 16.50% |
| MERS_CoV_B | NC_034440 | Bat | 16.08% |
| MERS_CoV_B | MG021451 | Bat | 14.91% |
| MERS_CoV_B | MG596802 | Bat | 15.89% |
| MERS_CoV_B | MG596803 | Bat | 15.75% |

TABLE S3

CORONAVIRUS SEQUENCES USED FOR MFED GENOME SCANS AND CONTTOUR PLOTS

| **Accession_no** | **Group** | **Isolate** |
|---|---|---|
| MN988713 | SARS-CoV-2 | Severe acute respiratory syndrome coronavirus 2 isolate |
| MT093571 | SARS-CoV-2 | Severe acute respiratory syndrome coronavirus 2 isolate |
| MT049951 | SARS-CoV-2 | Severe acute respiratory syndrome coronavirus 2 isolate |
| MT039890 | SARS-CoV-2 | Severe acute respiratory syndrome coronavirus 2 isolate SNU01 |
| MT027064 | SARS-CoV-2 | Severe acute respiratory syndrome coronavirus 2 isolate |
| MT007544 | SARS-CoV-2 | Severe acute respiratory syndrome coronavirus 2 isolate |
| MN994467 | SARS-CoV-2 | Severe acute respiratory syndrome coronavirus 2 isolate |
| MN996528 | SARS-CoV-2 | Severe acute respiratory syndrome coronavirus 2 isolate WIV04 |
| MN996527 | SARS-CoV-2 | Severe acute respiratory syndrome coronavirus 2 isolate WIV02 |
| FJ882953 | SARS-CoV-1 | SARS coronavirus MA15 ExoN1 isolate P3pp4 |
| AY654624 | SARS-CoV-1 | SARS coronavirus TJF |
| FJ882926 | SARS-CoV-1 | SARS coronavirus ExoN1 |
| FJ882943 | SARS-CoV-1 | SARS coronavirus MA15 ExoN1 |
| HQ890531 | SARS-CoV-1 | SARS coronavirus MA15 ExoN1 isolate d4ym1 |
| KF294457 | Bat sarbecovirus* | SARS-related bat coronavirus isolate Longquan-140 |
| GQ153543 | Bat sarbecovirus* | Bat SARS coronavirus HKU3-8 |
| GQ153547 | Bat sarbecovirus* | Bat SARS coronavirus HKU3-12 |
| DQ084200 | Bat sarbecovirus* | bat SARS coronavirus HKU3-3 |
| KJ473813 | Bat sarbecovirus* | BtRf-BetaCoV/SX2013 |
| KY770860 | Bat sarbecovirus* | Bat coronavirus isolate Jiyuan-84 |
| KJ473812 | Bat sarbecovirus* | BtRf-BetaCoV/HeB2013 |
| KJ473811 | Bat sarbecovirus* | BtRf-BetaCoV/JL2012 |
| KU182964 | Bat sarbecovirus* | Bat coronavirus isolate JTMC15 |
| KY938558 | Bat sarbecovirus* | Bat coronavirus strain 16BO133 |
| DQ648856 | Bat sarbecovirus* | Bat coronavirus (BtCoV/273/2005) |
| DQ412042 | Bat sarbecovirus* | Bat SARS coronavirus Rf1 |
| JX993987 | Bat sarbecovirus* | Bat coronavirus Rp/Shaanxi2011 |
| KP886809 | Bat sarbecovirus* | Bat SARS-like coronavirus YNLF_34C |
| DQ071615 | Bat sarbecovirus* | Bat SARS coronavirus Rp3 |
| KY417143 | Bat sarbecovirus* | Bat SARS-like coronavirus isolate Rs4081 |
| MK211377 | Bat sarbecovirus* | Coronavirus BtRs-BetaCoV/YN2018C |
| KY770858 | Bat sarbecovirus* | Bat coronavirus isolate Anlong-103 |
| KJ473816 | Bat sarbecovirus* | BtRs-BetaCoV/YN2013 |
| KY417145 | Bat sarbecovirus* | Bat SARS-like coronavirus isolate Rf4092 |
| FJ588686 | Bat sarbecovirus* | Bat SARS CoV Rs672/2006 |
| KY417142 | Bat sarbecovirus* | Bat SARS-like coronavirus isolate As6526 |
| MK211375 | Bat sarbecovirus* | Coronavirus BtRs-BetaCoV/YN2018A |
| KY417147 | Bat sarbecovirus* | Bat SARS-like coronavirus isolate Rs4237 |
| KY417148 | Bat sarbecovirus* | Bat SARS-like coronavirus isolate Rs4247 |
| KJ473815 | Bat sarbecovirus* | BtRs-BetaCoV/GX2013 |
| KY417146 | Bat sarbecovirus* | Bat SARS-like coronavirus isolate Rs4231 |

| | | |
|---|---|---|
| KC881006 | Bat sarbecovirus* | Bat SARS-like coronavirus Rs3367 |
| KC881005 | Bat sarbecovirus* | Bat SARS-like coronavirus RsSHC014 |
| KJ473814 | Bat sarbecovirus* | BtRs-BetaCoV/HuB2013 |
| DQ648857 | Bat sarbecovirus* | Bat coronavirus (BtCoV/279/2005) |
| DQ412043 | Bat sarbecovirus* | Bat SARS coronavirus Rm1 |
| MG772934 | Bat sarbecovirus* | Bat SARS-like coronavirus isolate bat-SL-CoVZXC21 |
| MG772933 | Bat sarbecovirus* | Bat SARS-like coronavirus isolate bat-SL-CoVZC45 |
| KY352407 | Bat sarbecovirus* | Severe acute respiratory syndrome-related coronavirus strain |
| GU190215 | Bat sarbecovirus* | Bat coronavirus BM48-31/BGR/2008 |
| JX993988 | Bat sarbecovirus* | Bat coronavirus Cp/Yunnan2011 |
| KF569996 | Bat sarbecovirus* | Rhinolophus affinis coronavirus isolate LYRa11 |
| MK211374 | Bat sarbecovirus* | Coronavirus BtRl-BetaCoV/SC2018 |

*Used for contour plots only

TABLE S4

PREDICTED RNA STRUCTURE ELEMENTS IN CORONAVIRUS GENOMES

| Virus | Duplexes | Max. length | Stem-loops | Length range | Prop. paired |
|---|---|---|---|---|---|
| SARS-CoV-2 | 2015 | 14 | 657 | 3-44 | 63.0% |
| SARS-CoV | 1934 | 14 | 645 | 3-39 | 65.7% |
| MERS-CoV | 2003 | 16 | 627 | 3-18 | 62.3% |
| OC43 | 2034 | 18 | 616 | 3-24 | 66.9% |
| HKU1 | 1877 | 16 | 557 | 3-21 | 65.7% |
| 229E | 1909 | 15 | 473 | 3-16 | 66.5% |
| NL63 | 1798 | 17 | 500 | 3-18 | 65.6% |
| Deltacoronavirus | 1763 | 14 | 547 | 3-18 | 64.2% |
